# Prediction and characterization of transcription factors involved in drought stress response

**DOI:** 10.1101/2020.04.29.068379

**Authors:** Chirag Gupta, Venkategowda Ramegowda, Supratim Basu, Andy Pereira

## Abstract

Transcription factors (TFs) play a central role in regulating molecular level responses of plants to external stresses such as water limiting conditions, but identification of such TFs in the genome remains a challenge. Here, we describe a network-based supervised machine learning framework that accurately predicts and ranks all TFs in the genome according to their potential association with drought tolerance. We show that top ranked regulators fall mainly into two ‘age’ groups; genes that appeared first in land plants and genes that emerged later in the *Oryza* clade. TFs predicted to be high in the ranking belong to specific gene families, have relatively simple intron/exon and protein structures, and functionally converge to regulate primary and secondary metabolism pathways. Repeated trials of nested cross-validation tests showed that models trained only on regulatory network patterns, inferred from large transcriptome datasets, outperform models trained on heterogenous genomic features in the prediction of known drought response regulators. A new R/Shiny based web application, called the DroughtApp, provides a primer for generation of new testable hypotheses related to regulation of drought stress response. Furthermore, to test the system we experimentally validated predictions on the functional role of the rice transcription factor *OsbHLH148*, using RNA sequencing of knockout mutants in response to drought stress and protein-DNA interaction assays. Our study exemplifies the integration of domain knowledge for prioritization of regulatory genes in biological pathways of well-studied agricultural traits.

**One Sentence Summary:** Network-based supervised machine learning accurately predicts transcription factors involved in drought tolerance.

---

The drastic reduction in soil water content negatively regulates growth and development of crop plants such as rice (*Oryza sativa*), causing substantial loss in yield and quality (Boyer, 1982; Bray, 1997; Yamaguchi-Shinozaki and Shinozaki, 2006; Palanog et al., 2014). Plants and specific genotypes within a plant species that can withstand reduced soil water content would be identified as ‘drought tolerant’, and offer examples to study the mechanisms involved in their survival and productivity in terms of yield. While conventional breeding has been the preferred method of improving drought tolerance in rice and other crop plants, modern genomics and genetic engineering strategies have become integral part of trait enhancement programs (Umezawa et al., 2006; Ashraf, 2010; Gaj et al., 2013). A prerequisite for effective use of genetic engineering tools in trait improvement is the prior knowledge about candidate genes that are likely to produce a desirable phenotype when genetically intervened. Although transcriptome analysis of rice under water limited conditions has identified thousands of differentially expressed genes, it is difficult to narrow down the selection of candidate genes for testing function and genetic modification. This lack of candidate genes will be a major bottleneck in future, as it impedes our ability to set up targeted genetic screens to select leads for further crop improvement (Gutterson and Zhang, 2004; Century et al., 2008; Jansing et al., 2019; Baxter, 2020). Therefore, new versatile computational methods and data-driven approaches capable of discovering key genes regulating complex traits like drought tolerance are needed.

Gene regulatory networks (GRN) play a central role in mediating plant responses to environmental stresses (Chen and Zhu, 2004; Clauw et al., 2016; Lovell et al., 2018). Transcription Factors (TFs) are key nodes (genes) in these networks as they regulate the expression of several downstream genes involved in many stress responsive pathways and biological processes (Yang et al., 2011). Therefore, TFs remain the most appealing candidates for genetic engineering of stress tolerance due to their regulatory nature (Tran et al., 2010; Rabara et al., 2014; Krannich et al., 2015; Wang et al., 2016; Hoang et al., 2017). Computational modeling of genome-scale regulatory networks inferred from large-scale transcriptomic datasets is a feasible approach (Razaghi-Moghadam and Nikoloski, 2020), and has shown great promise in accelerating the process of *in silico* gene discovery to *in planta* gene validation. Some good examples of recent plant studies that used large-scale GRNs to discover novel gene functions are outlined in recent review articles (Li et al., 2015; Gupta and Pereira, 2019; Haque et al., 2019).

There are several limitations of most of the popular approaches currently used to mine relevant biological signals from network data. For example, function interpretation of network neighborhoods (modules, clusters etc.) in terms of known biological processes and pathways is only secondary knowledge, which does not directly allow either module or gene prioritization, that can also be objectively tested. Also, gene groups that remain unannotated cannot simply be used in interpretation of the inferred network model. Furthermore, the concept of ‘hub’ genes in a context-specific network has limited interpretation and cannot be generalized across network-types (Langfelder et al., 2013; Walley et al., 2016; Vandereyken et al., 2018). Newer computational approaches for candidate gene prioritization are needed (Liseron-Monfils et al., 2018; Dursun et al., 2019), including methods that allow gene prioritization informed by integrating networks with new experimental data such as GWAS results (Schaefer et al., 2018). We also need to develop methods that can leverage on prior documented knowledge about gene-phenotype and gene-trait links to make genome-wide predictions, for example, when a dedicated GWAS study for the trait is unavailable.

In rice, the function of a few TFs involved in multiple responses to water-deficit conditions have been identified by overexpression or loss-of-function analysis. These experimentally validated ‘gold-standard’ examples of drought regulators provide an opportunity to test the feasibility of generating machine learning models predictive of other untested drought TFs. Recently, supervised machine learning has been very useful in generation of predictive models for various aspects of research in plant and crop biology (Ma et al., 2014; Sperschneider, 2019). For trait-gene predictions, binary classifiers – algorithms that classify genes into two classes based on their discriminative attributes– seem to be very popular among plant biologists. For example, thousands of genomic and evolutionary features that characterize known essential genes were used to train models predictive of other lethal-phenotype genes (Lloyd et al., 2015). Similarly, several distinguishing features of genes currently annotated in secondary or primary metabolism pathways were used to train models capable of predicting new specialized metabolism genes (Moore et al., 2019). Putative cis-regulatory elements (CREs) involved in general abiotic and biotic stress responses (Zou et al., 2011), and CREs involved in regulation of root cell type responses to high salinity stress (Uygun et al., 2019) have also been identified by training supervised learning models. Particularly interesting are the studies that used an inferred genome-scale network, instead of heterogenous genomic features, as input to the learning algorithm to make reliable genome-wide predictions on disease-gene associations in human (Guan et al., 2010; Guan et al., 2012; Krishnan et al., 2016; Liu et al., 2019). However, whether this network-based supervised machine learning approach can be applied to predict TFs associated with certain traits of interest still remains to be tested.

To determine the feasibility of predicting TFs likely involved in drought tolerance (DT) mechanisms in rice, we first inferred the consensus GRN in response to abiotic-stress response using an ensemble of reverse-engineering algorithms. We leveraged on documented phenotypes associated with rice TFs listed in multiple databases, and trained machine learning models that learnt regulatory network patterns characteristic of drought response. Application of the trained model resulted in predictions where all TFs in the genome were scored along a continuous spectrum according to their potential association to DT. We then described the phylostratigraphic, structural and functional features of TFs at both ends of this spectrum. Finally, we tested the effect of using 1) only the consensus GRN, 2) only newly inferred genomic features and 3) integration of the network and genomic features on overall accuracy of the models in predicting known drought response TFs kept hidden (hold-out set) in the training process **(Fig. 1)**. These features of TFs that likely regulate drought stress responses will be important in gene prioritization for experimental validation and genetic enhancement of drought tolerance in rice.

**Figure 1:**
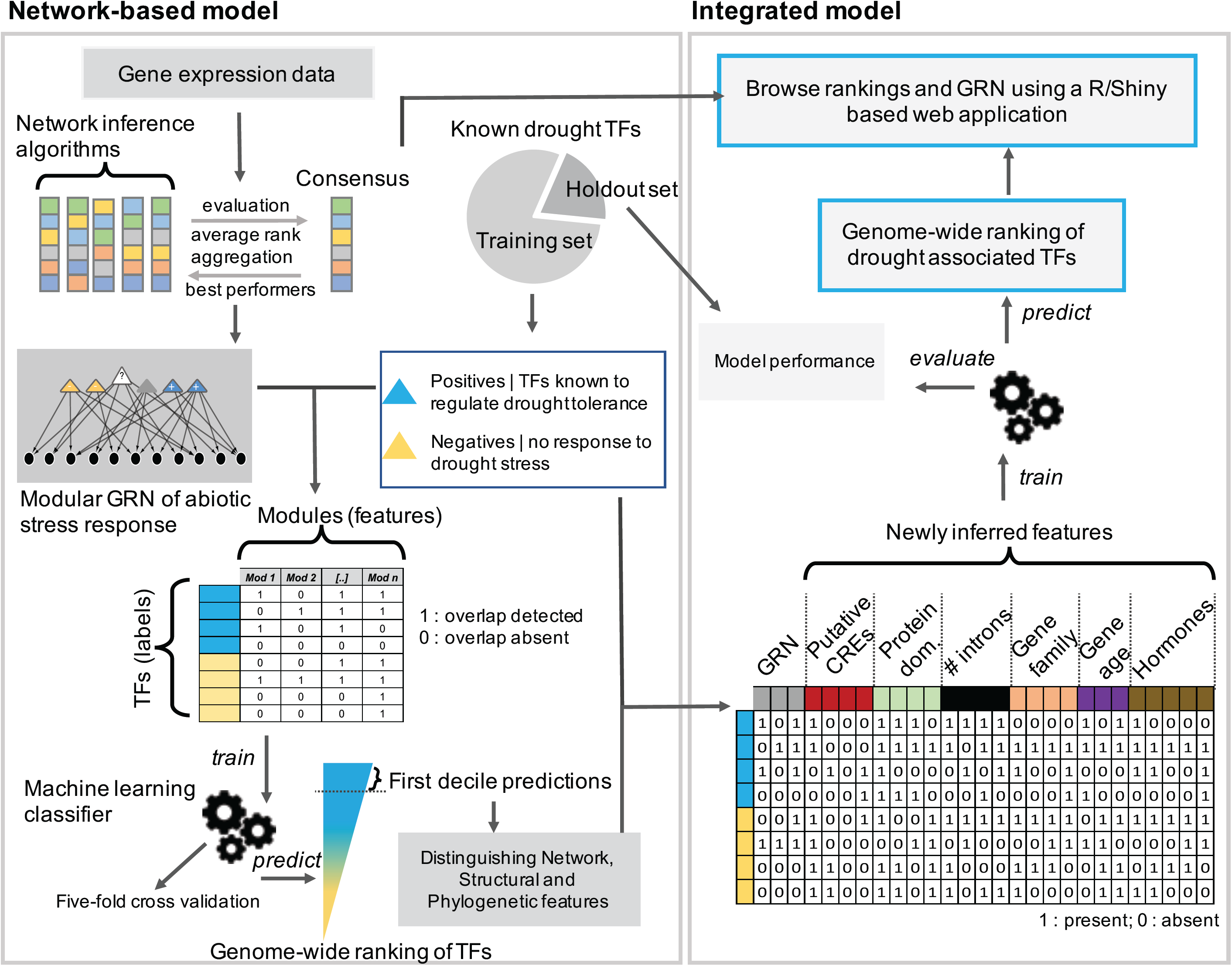
Workflow of the network-based machine learning approach used in this study. A consensus modular gene regulatory network (GRN), representing relationships between TFs and functional modules, was predicted from expression data using an ensemble of network prediction algorithms. Rice knowledgebases were mined to identify TFs that are already reported as regulators of drought tolerance (labeled as drought positive class), and a set of TFs that did not respond to drought in published gene expression studies (labeled as drought negative class). These benchmark drought TFs, along with their network connectivity patterns in the consensus GRN, were used as input training data for a binary classification algorithm (support vector machine) to identify patterns that can discriminate between the two classes of benchmark TFs. The identified patterns were subsequently used to classify the remaining unlabeled TFs. The final output of this supervised network-based model was the representation of each TF in the rice genome along a continuous spectrum representing its association to drought tolerance. Discriminative genomic features of TFs at both the ends of this spectrum were identified and described. These newly inferred genomic features were then integrated with the network-based features and evaluated for accuracy using nested cross-validation tests, where the outer loop was a two-fold split and the inner loop was a five-fold split. The GRN and predictions can be accessed online at http://rrn.uark.edu/shiny/apps/rrn/.

## Results and Discussion

### Inference of the consensus modular gene regulatory network in response to global abiotic stress response

We started with inference of the global gene regulatory network (GRN), from a large collection of publicly available gene expression datasets conditioned on abiotic stress responses. Instead of relying on any one of the several competing algorithms frequently used for inference of GRNs, we created an ensemble of five complementary methods and statistically aggregated the outputs of these methods to create a consensus GRN (Marbach et al., 2012). The aggregation of networks inferred from different algorithms was necessary as they individually showed very little overlap between inferred TF-gene links (**Fig. 2A; Supplemental Data S1)**.

**Figure 2:**
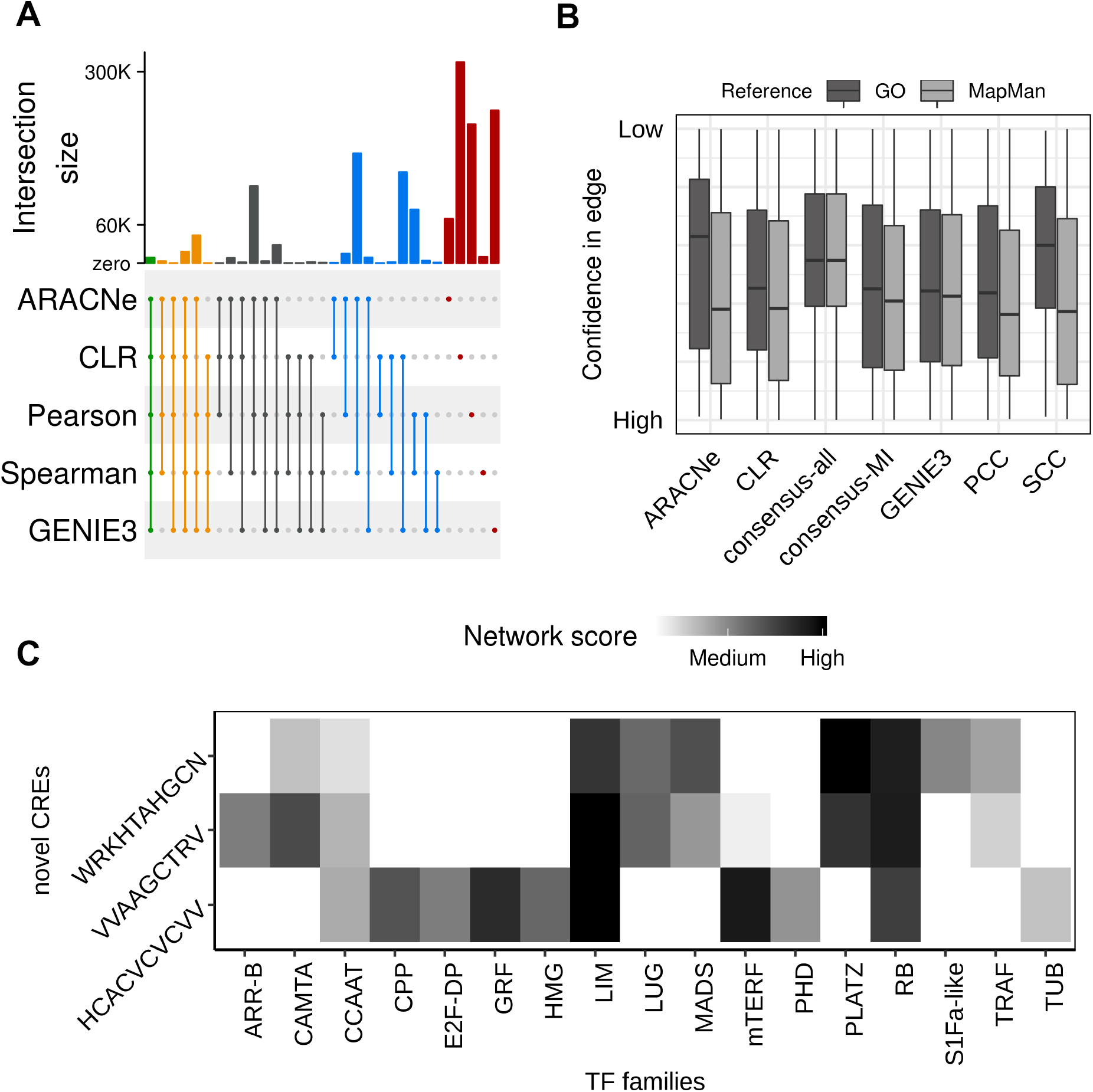
Inference, evaluation and functional annotation of the rice gene regulatory network. A) An upset plot showing the overlap between edges predicted by different network prediction algorithms and their aggregate. The bars on the top indicate the size of overlap between methods connected by dots in the center matrix. Each intersection is color coded uniquely. Red: Unique edges, Blue: Overlap between two methods, Black: three methods, Orange: four methods, Green: all five methods. B) The boxplots show the distribution of ranks given to ‘reference’ edges derived from the Gene Ontology (GO) and MapMan pathways by each network prediction method in the ensemble. Consensus-all indicates an aggregate solution of all five methods and consensus-MI indicates an aggregate solution of only mutual-information-based methods. C) Besides recovering several known plant CREs, the *de novo* CRE analysis identified three novel motifs that did not match to any known plant CRE listed in multiple databases. The heatmap shows that these novel motifs could potentially be direct or ‘associative’ binding sites of members from seven TF families, based on significant overlaps of the predicted targets of TFs from the families on the *x* axis within the genes that harbor the three novel CREs on the *y* axis (hypergeometric tests *q* value < 0.01). Color gradient indicates the network score, calculated as the average ranks of edges from the consensus gene regulatory network. Darker color indicates stronger association between the CRE and the TF family, as indicated in the key.

We evaluated the performance of each algorithm in our ensemble, based on their ability in correctly predicting 1) genes linked with position weight matrices of rice TFs listed in the CIS-BP database (Weirauch et al., 2014), and 2) co-annotated TF-gene pairs from specific biological process categories from the latest version of rice Gene Ontology (GO) (Ashburner et al., 2000) as well as pathway annotation bins in rice from MapMan (Thimm et al., 2004). Both these evaluations showed that aggregating outputs from different network prediction algorithms was generally better in terms of accuracy (estimated as an *F*-score; Supplemental Table 1). This evaluation also showed that methods that use mutual information as a base measure to capture direct functional relationships between TFs and potential target genes performed better than the simple correlation-based methods **(Fig. 2B)**. Therefore, the aggregate of three mutual information methods was chosen as the consensus GRN and used in further analysis.

We next computed the level of ‘coregulation’ between functional genes in the inferred GRN (see *Methods)*, and applied a network clustering algorithm to group genes into modules of highly coregulated genes (van Dongen and Abreu-Goodger, 2012) (**Supplemental Data S2)**. Out of the 740 modules thus obtained, the biological relevance of 31% could be verified using enrichment analysis of function annotation data from various sources. Additionally, ∼41% of all modules were found preserved in an independent coexpression network built earlier (Krishnan et al., 2017). Interestingly, 22% of these preserved modules were found amongst the ones that could not be annotated by gleaning function annotation databases, indicating that these are biologically relevant gene groupings that fill large gaps that still exist in the current state of functional annotations in rice (**Supplemental Data S3**). In addition to function enrichment to annotate modules, we also performed a *de novo* analysis of *cis* regulatory elements (CREs) in the promoter regions of the module genes (Elemento et al., 2007). We expected this *de novo* analysis to recover known and novel <u>a</u>biotic <u>s</u>tress related CREs (AS-CRE), given the context of the underlying network. The analysis detected a total of 84 AS-CREs distributed across modules, and 81 of these AS-CREs matched to putative CREs listed in multiple plant databases (**Fig. S1A and S1B; Supplemental Data S4;** see *Additional notes*). Interestingly, network analysis of these CREs indicated that two of the three unmatched novel DNA motifs could likely be binding sites of TFs from the same families **(Fig. 2C)**.

Because function enrichment analysis and the analysis of AS-CREs showed that most predicted modules constitute biologically relevant gene groupings, we assigned TFs as potential regulators of modules based on the overlap (estimated using Jaccard’s similarity) between genes within each module and the predicted targets of each TF in the consensus GRN (see *Additional notes*). As illustrated in **Figure 3A**, the inferred relationships between TFs and modules of coregulated genes is structured like a weighted matrix - with several layers of annotations on modules to allow biological interpretation - representing a global transcriptional regulatory map of abiotic stress responses in rice. An R/shiny-based application was also developed to allow browsing the network with a gene of interest though a web browser (http://rrn.uark.edu/shiny/apps/rrn/).

**Figure 3:**
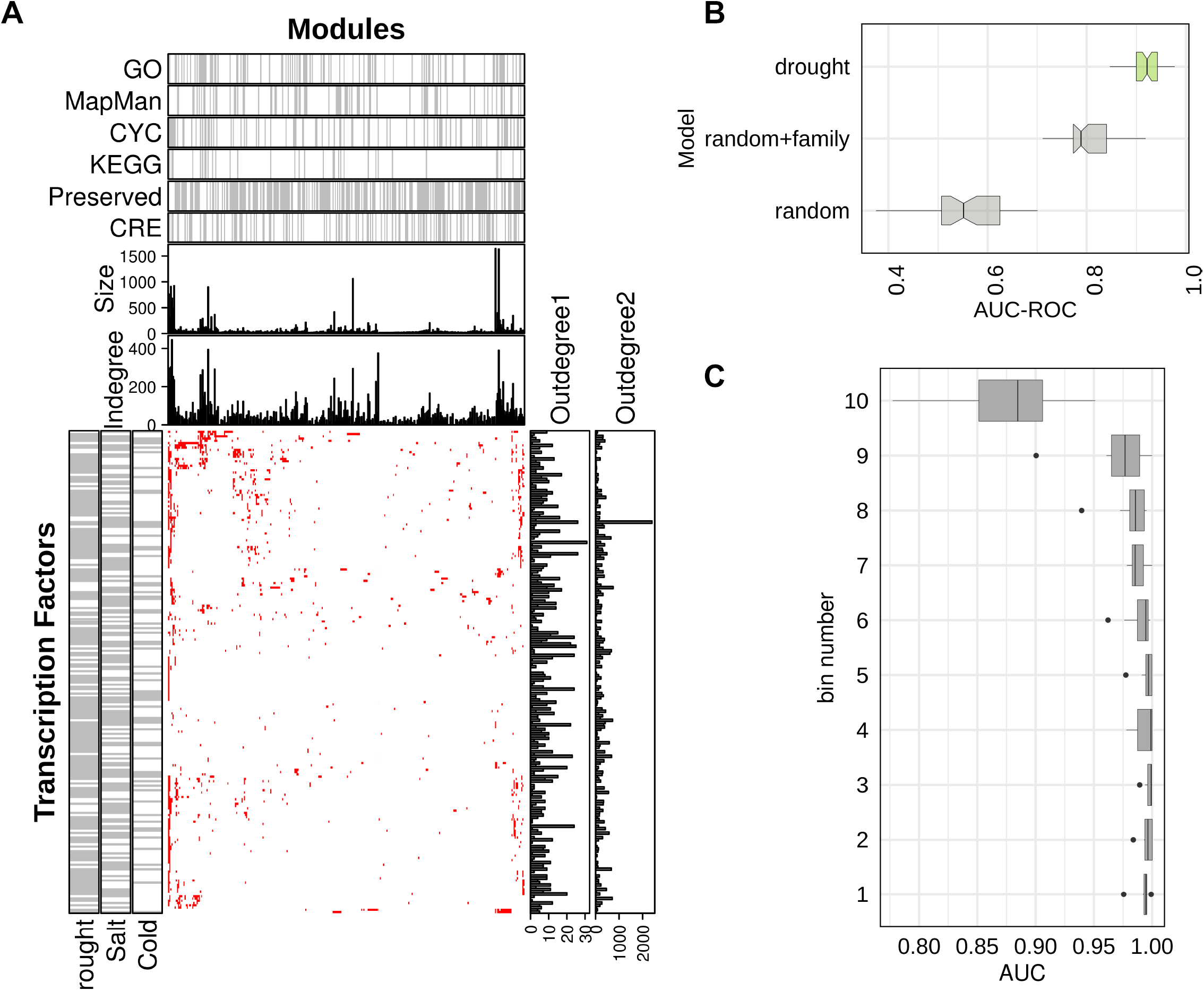
Evaluation of the network-based classifier. A) An annotated heatmap (bottom center) depicting modules (columns) along with their potential regulators (rows). The cells of the heatmap are colored red if an overlap of at least one gene was found between the predicted targets of the TF and genes in the module. Other cells are colored white. TFs reported to be involved in drought, salt and cold stress response are indicated by grey horizontal bars (bottom left). Outdegree 1 and 2 bar plots (bottom right) indicate number of genes and number of modules predicted to be targeted by each TF in the corresponding row. Module annotations are illustrated on the top of the heatmap. Indegree and size bar plots indicate the number of incoming edges and the size of each module, respectively. Modules significantly enriched with functional categories from four function annotation databases, preserved network modules and CREs are indicated by vertical grey bars (top). B) Boxplots showing the distribution of area under the receiver operator curve (AUC; *x* axis) of the classifier trained using reported drought tolerance genes (shaded green; top), the classifier trained using randomly picked TFs (bottom), and the classifier trained using randomly picked TFs with distribution of families equal to that of the drought classifier (center). C) TFs were sorted according to their decreasing drought scores and grouped into 100 equal-sized bins. Expression levels (transcript per million units) of TFs in each bin was used as features to classify a set of labeled RNA-seq samples as drought or control (GSE74793). Each boxplot shows distribution of AUC scores *(x* axis) from three-fold cross validation tests in groups of 10 bins, with lower numbered bins (*y* axis) indicating TFs with higher drought scores.

### Network-based supervised machine learning enables prediction of transcription factors involved in drought tolerance

While the consensus GRN we described above can potentially benefit gene function predictions using typical ‘gene-neighborhood’ analysis, we next demonstrate that this network can also be used in a machine learning framework for systematic genome-wide prioritization of TFs that likely regulate drought stress responses in rice. To generate the training data for supervised modeling, we surveyed the functional rice gene database (Yao et al., 2018), the rice mutant database (Zhang et al., 2006) and the Oryzabase (Kurata and Yamazaki, 2006), and retrieved all rice genes with documented phenotypes under drought or water-limiting conditions on the basis of experiments on loss-of-function mutants or transgenic overexpression lines. Because of the complex genetic basis of drought responses, this list of ‘drought associated’ genes do not represent any particular physiological, morphological or biochemical phenotype typically measured in the analysis of drought stress tolerance response. Therefore, we use ‘drought tolerance’ (DT) as a term to broadly encapsulate various definitions of ‘drought stress response’, representing global molecular mechanisms by which plants adapt, escape or otherwise respond to water limiting conditions (Basu et al., 2016). As of May 2019, we found 165 TFs amongst all the DT genes obtained from database mining. We labeled these TFs as the ‘drought positive’ class, and 682 TFs that did not respond to drought stress in reanalysis of a number of published gene expression datasets (and other public resources) as the ‘drought negative’ class (see *Methods)*. The remaining TFs not found in any of these two classes were left unlabeled **(Supplemental Data S5)**.

The problem of DT TF prediction was then formulated as a two-class classification problem, where the goal was to predict the class label of each unlabeled TF. To achieve this, the support vector machine (SVM), a binary classification algorithm, was used to train models that learnt regulatory network patterns discriminative of the drought positive and negative classes of TFs. The accuracy of trained models was evaluated using five-fold cross validation tests. This test splits all training examples (drought positive and negative TFs) into five equal parts. The model is trained on four of the five splits and tested on the remaining split kept hidden in training, ensuring that each split is used as the test-set only once. The accuracy of the model was evaluated using <u>a</u>rea <u>u</u>nder the receiver-operator <u>c</u>urve (AUC) statistics. The AUC ranges between 0 and 1, with values closer to 1 indicating superior performance in classifying test-set TFs in their respective class. In 10 independent runs of five-fold cross validation tests, our network-based DT classifier achieved an average AUC of 0.91, which is significantly larger than the model trained using randomly picked TFs from the genome **(Fig. 3B)**. The observed AUC of the DT classifier was also found to be significantly better than the model trained using randomly picked positives, while maintaining family memberships and class size distributions similar to that of the real positive examples **(Fig. 3B)**. This indicated that even if TFs within the same family are more likely to be functionally similar, this cannot be the only deterministic feature for classification.

Application of the validated model to the whole dataset (including other unlabeled TFs) resulted in a rank for each TF. To ease interpretation, we scaled these ranks between 0 and 1 such that a threshold of 0.9 meant TFs in the top 10% predictions, 0.8 meant top 20% predictions and so forth. Therefore, this technique placed 2160 TFs (> 93% of all known TFs in rice) along a continuous spectrum of drought scores (DS) representing their potential association to DT (**Supplemental Data S5**).

### Occurrence of drought can be accurately inferred from the expression levels of TFs with high drought scores

We next evaluated the DS produced by the network-based classifier described above by reanalyzing a recently published RNA-seq dataset of rice seedlings exposed to drought (Wilkins et al., 2016). We hypothesized that if TFs with larger DS are true regulators of drought, their expression levels should be indicative of whether the plant has sensed drought or not. To test this, we first divided all TFs into 100 bins based on decreasing DS, with each bin consisting of ∼21 TFs (total 2160 TFs). Therefore bin #1 consisted of top 1% predictions, bin #2 consisted of top 2-3% predictions, bin #3 consisted of top 3-4% predictions, and so forth. We then evaluated whether expression levels of TFs in each bin can correctly classify a sample in the seedling RNA-seq as drought or control. We observed that bins with larger DS scores are generally more accurate in this classification as compared to bins containing TFs with smaller DS (**Fig. 3C)**. This indicated that the network-based classifier placed potentially true regulators of DT toward the top of the rankings. Hence, the prioritization is correct and TFs at the very top of the rankings are reliable candidates for characterization of regulatory mechanisms involved in DT.

### The drought score correlates with known expression patterns and high scores are indicative of conserved responses

Because these evaluations suggested that our approach of prioritizing TFs involved in DT is reliable, we asked if the TFs that are at the top of the rankings have specific characteristics that can distinguish them from TFs at the bottom of the rankings. We began our initial investigations by first examining relationships between predicted DS of TFs and their expression patterns in various contexts. Using the spatial and temporal drought response dataset (Wang et al., 2011), and our previous developmental stage drought response dataset (Krishnan et al., 2017), we found that differentially expressed (t-test *q* value < 0.05) TFs in both these datasets have significantly larger (Welch’s t-test *p* value < 0.001) mean DS compared to the mean DS of the background of remaining TFs that did not differentially express **(Fig. 4A and 4B)**. Next, since phytohormones are known to mediate drought stress responses (Kazan, 2015; Muller and Munne-Bosch, 2015; Sah et al., 2016; Ullah et al., 2018) and also modulate activities of TFs (Liu et al., 2012; Banerjee and Roychoudhury, 2017), we examined the hormone-exposed seedling dataset (Garg et al., 2012). TFs that differentially expressed in response to six phytohormones in this hormone dataset have significantly larger mean DS (range 0.62-0.79) than that of TFs that did not respond **(Fig. 4C)** (*p* values < 1.12e-06). These datasets established a positive relationship between predicted DS and known expression patterns of rice TFs.

**Figure 4:**
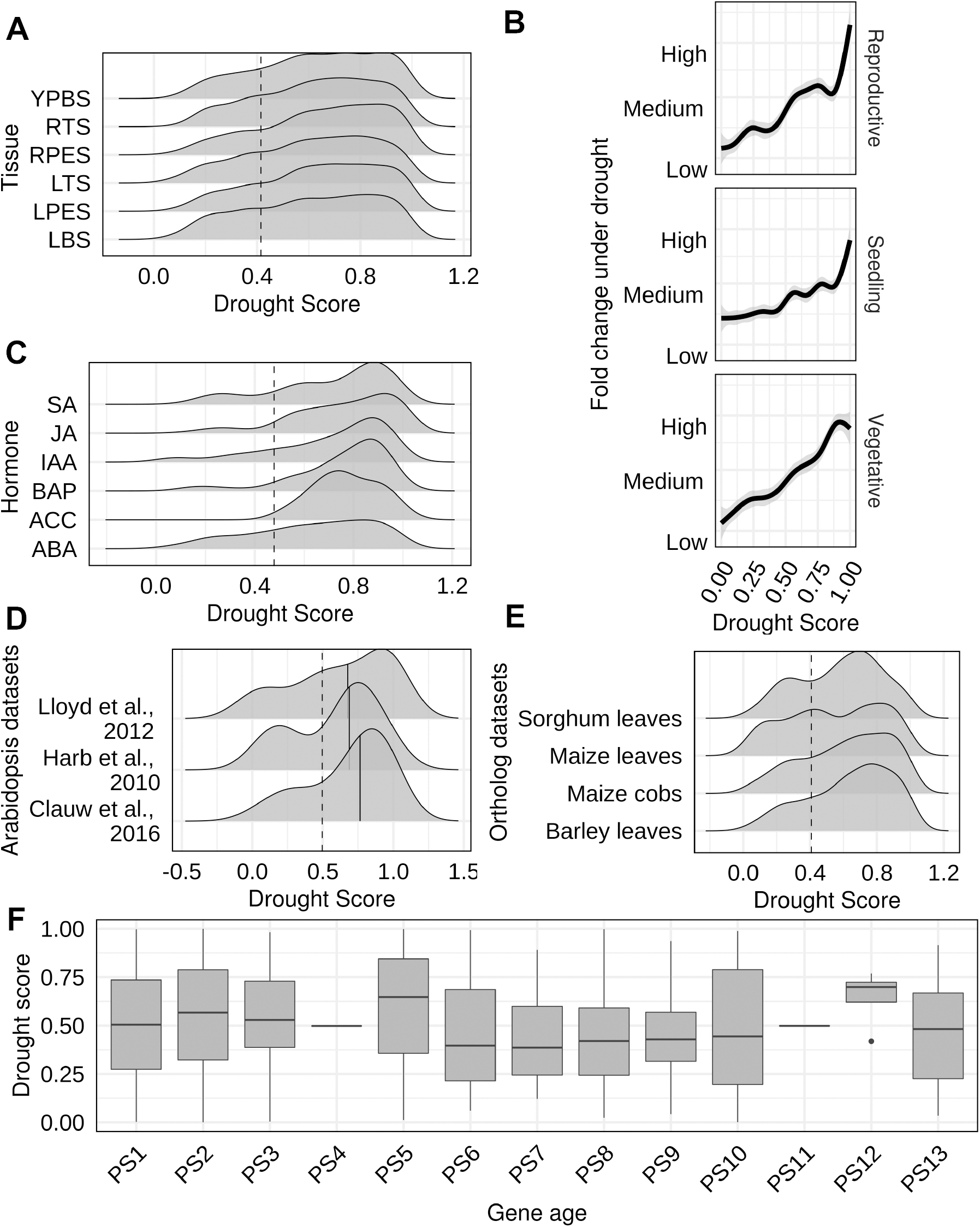
Relationships between drought scores, ortholog gene expression and phylostratigraphic profiles of rice TFs. TFs that differentially expressed in A) spatial and temporal drought response dataset (GSE26280), B) three different stages of development in the reference genome (Nipponbare; MSU7) and C) response to hormone treatments have significantly higher drought scores (DS) compared with DS of the background of all remaining TFs in each case. (YPBS: young panicle booting stage, RTS: root at tillering stage, RPES: roots at panicle elongation stage, LTS: leaves at tillering stage, LPES: leaves at panicle elongation stage, LBS: leaves at booting stage; IAA: Indole-3-acetic acid (auxin), BAP: benzyl aminopurine (cytokinin), ABA: abscisic acid, ACC: 1-aminocyclopropane-1-carboxylic acid (ethylene derivative), SA: salicylic acid, JA: jasmonic acid). D) Rice TFs with orthologs in Arabidopsis genes that differentially express under drought stress (center), or are known by experimental validation (top) or predicted for drought tolerance (bottom) have significantly higher DS compared with the background. E) Similarly, DS of TFs (*x* axis) with orthologs in genes that differentially expressed in different crop datasets (*y* axis) is also skewed toward larger DS values. F) Box plot showing the distribution of DS in different age groups according to NCBI taxonomic classification. PS5 and PS12 represent Embryophytes and Oryza clades, respectively.

To examine whether TFs with high DS also have conserved expression patterns, we examined the drought response of orthologous rice genes in datasets from other plants. Along with a set of differentially expressed Arabidopsis genes that responded to mild and severe drought assays we reported previously (Harb et al., 2010), we included a set of drought genes with experimental evidence listed in the Arabidopsis phenotype database (Lloyd and Meinke, 2012), and a set of Arabidopsis genes recently predicted to be involved in mild drought responses (Clauw et al., 2016). As illustrated in **Figure 4D**, the distributions of rice TFs with orthologs in these three sets were found skewed toward larger values of DS. The mean DS of these orthologous rice TF ranges between 0.57-0.68, which is significantly larger than the mean DS of 0.49 of the background of remaining 2106 TFs (*p* value = 0.00068). This indicated that rice TFs with larger DS have conserved drought responses in Arabidopsis.

Furthermore, such skewed distributions were also observed in reanalysis of drought response datasets from other cereal crops **(Fig. 4E)**; rice TFs with orthologs that differentially expressed in response to application of drought stress in 1) cobs and 2) leaves of maize (Kakumanu et al., 2012), 3) leaves of leaves of barley (Cantalapiedra et al., 2017), and 4) leaves of sorghum have significantly larger mean DS (*p* values < 0.05) than the mean of the background in all cases **(Fig. 4E**). Interestingly, the distribution of DS was observed to be bimodal and fell more towards the middle in the leaves of maize and sorghum, although the mean DS was weakly but significantly larger than the background (*p* value=0.02). These differences specifically in the leaves of maize and sorghum could be due to differences in their mode of photosynthesis compared to rice and barely. Further testing with data from drought exposed leaves and non-photosynthetic tissues of other C3 and C4 crops is needed to build testable hypotheses around predictable drought-photosynthesis relationships from the network. Overall, these datasets suggest that predicted DT TFs could be functionally conserved for responses to drought stress in other plants and crops.

We next asked if the predicted DS and evolutionary age of a TF are related. Using phylostratigraphic profiles of rice genes (Wang et al., 2018), we observed two peaks in DS within the 13 phylostratum (PS) age groups rice genes fall into. The first peak in PS5, which corresponds with the Embryophytes (land plants) clade and the second peak in PS12, which coincides with Oryza clade, both mirror major events in evolutionary history of rice **(Fig. 4F)**. To examine the distribution of DS of TFs that arose in the terminal clade (*O. sativa*, closely related rice varieties), we examined the available pan genome of rice (Sun et al., 2017), but did not find any significant differences in DS between core and distributed TFs, or TFs that are *indica*- or *japonica*-dominant (**Fig. S2A-C**). This analysis indicated that most of the top ranked TFs are conserved across land plants, while the youngest TFs with high DS could have evolved specifically during adaptation of rice to drought. While the functions of relatively younger genes still remains difficult to predict from expression data alone (Ruprecht et al., 2017; Hansen et al., 2018), it would be interesting to explore their roles in drought response from the lens of our analysis.

### Structural characteristics of predicted drought tolerance transcription factors

Recent studies in rice and other organisms suggest that younger genes have relatively simple exon/intron and protein structure (Neme and Tautz, 2013; Cui et al., 2015; Wang et al., 2018). Other studies have showed that simple genes, for example those that lack introns, are rapidly regulated (Jeffares et al., 2008; Speth et al., 2018), and such genes represent an important component of the possibly conserved stress response machinery in land plants (Jeffares et al., 2008; Zhu et al., 2016; Morozov and Solovyev, 2019). Since most of the TFs strongly predicted to be associated with DT in our analysis are also the ones that first emerged in land plants, we next investigated if the structural features of TFs at the top of our rankings also have simple gene-body structure and protein domain features. Indeed, the top ranked TFs were found to be generally intron-poor genes, and a significantly large proportion (chi-square test *p* value=0.0058) of them are intronless (**Fig. 5A**). However, the coding sequence length of TFs at the top of the rankings was not different from the background of all remaining TFs, but significantly larger than TFs at the bottom of the rankings (*p* value=5.616e-11) **(Fig. 5B)**. In addition, ∼75% of top ranked TFs were predicted to encode small proteins with either one or two InterPro domain annotations (**Fig. 5C)**. We also confirmed that the classifier did not assign high DS to known pseudogenes in rice (Karro et al.; Karro et al., 2006; Thibaud-Nissen et al., 2009), indicating that even if they emerged due to loss of protein domain(s) in a parent gene, their function was either retained or altered to benefit the species (**Fig. S3**).

**Figure 5:**
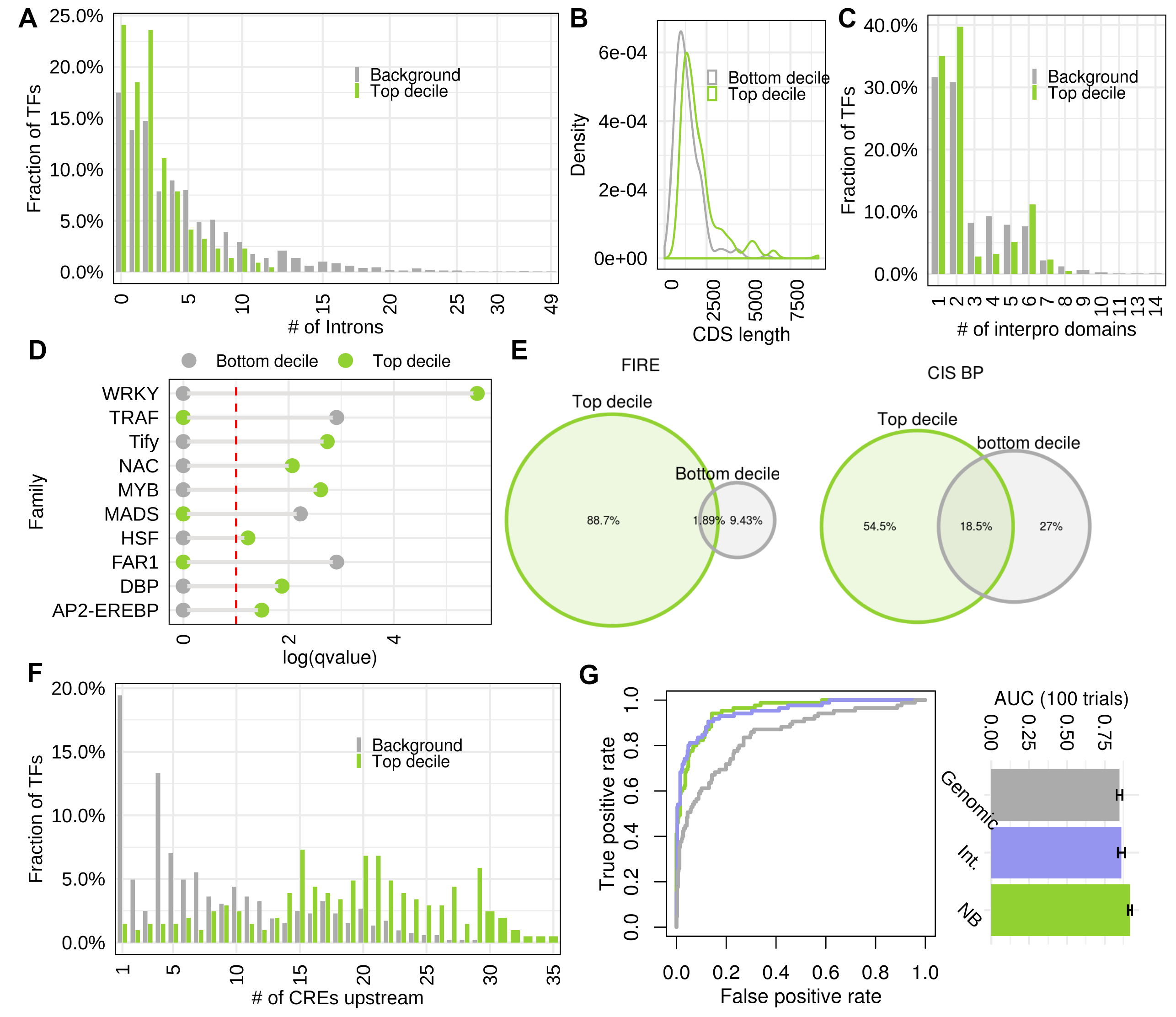
Structural features of predicted drought tolerance transcription factors and their evaluation. The first decile TFs (top 10% predictions) are A) intron poor compared to background of remaining 90% TFs, and this pattern continues till top 40% predictions. B) However, no significant differences between the coding sequence length of TFs at the top and bottom of the rankings was observed. C) Top 20% predictions contain fewer protein domains compared to the background. D) A dumbbell plot showing enrichment of TF families within the top decile (green dots) and bottom decile (grey dots). The -log of Storey’s *q* values resulting from hypergeometric tests is represented along the *x* axis and the families indicated along the *y* axis. E) Venn diagrams showing low overlaps between DNA binding motifs linked with top and bottom decile TFs. Left panel shows motifs identified from *de novo* analysis (using FIRE; see supplemental methods) and the right panel shows motifs listed in the CIS-BP database. F) Top decile predictions have a larger number of CREs present within 1000 bp upstream promoters compared to the background. G) All these genomic features alone are less accurate in correctly predicting known regulators of drought, as shown by the receiver operator curve (grey line, left panel) compared to the classifier that used only network-based features (green line), and the classifier trained by integrating genomic and network features (blue line). The network-based (NB) model performed with highest average accuracy in 100 random trails of nested cross-validation tests (bar plot right panel).

In terms of gene families, the top ranked drought TFs were found enriched with WRKY, Tify, NAC, MYB and AP2/ERF families (FDR corrected hypergeometric test *p* values < 0.1) (**Fig. 5D)**. These gene families are well-known to associate with drought stress in multiple crops (Yu et al., 2012; Gahlaut et al., 2016; Hoang et al., 2017). In contrast, TFs at the bottom of the rankings were found enriched in growth and development associated gene families such as the MADS, FAR1 and TRAF (Smaczniak et al., 2012; Tedeschi et al., 2017; Ma and Li, 2018). We also observed that the TFs at the top and bottom of the rankings likely bind to distinct groups of DNA binding sites **(Fig. 5E)**, and the *de novo* predicted AS-CREs from network modules occur more frequently within the promoters of top ranked TFs **(Fig. 5F)**, indicating presence of a hierarchical response system. Overall, this analysis indicated that the drought classifier clearly discriminated between features of stress and development-related gene families, although this information was not explicitly encoded in the input set of features used to train the model.

### Network-based learning outperforms learning from genomic features

The network-based machine learning model described above revealed several interesting features of TFs that are likely involved in drought response mechanisms, and these inferred features generally agree with our current understanding about abiotic stress responses in plants. Therefore, we next tested the feasibility of training the DT classifier using only these inferred genomic features. We reasoned that if these features are truly discriminative, they should be predictive of known DT TFs. To perform an unbiased evaluation, 50% of all training labels (∼422 TFs) were randomly selected as a hold-out evaluation set, and the model was trained and cross-validated on the remaining 50%. We observed that at any given true positive rate threshold, the model trained using only the inferred genomic features, although better than random, consistently attained higher false positive rates compared to the model trained using only regulatory network patterns of TFs (**Fig. 5G, left)**. In 100 repeated trials, the average AUC score of the network-based model was found to be significantly higher than the genomic model (*p* value < 2.2e-16), as well as the model trained by fusing genomic features with the network. The network-based model was also more robust to variation in training labels relative to the other two models **(Fig. 5G, right)**.

### Predicted drought tolerance transcription factors are involved in hormone-mediated responses

Because the network-based machine learning model outperformed the model trained on genomic features, we finally investigated the network modules that served as best predictors of this classification to gain functional insights on predicted drought regulators. Using ‘feature importance’ scores from the model output, we found that 22% of all modules predicted DT positively (**Supplemental Data S6)**. We connected these drought modules with their predicted regulators, linked the AS-CREs to modules as well as TFs (using FDR corrected hypergeometric test *p* value threshold < 0.01), and explored this interconnected multi-node network in Cytoscape (Shannon et al., 2003) **(Fig. 6A)**.

**Figure 6.**
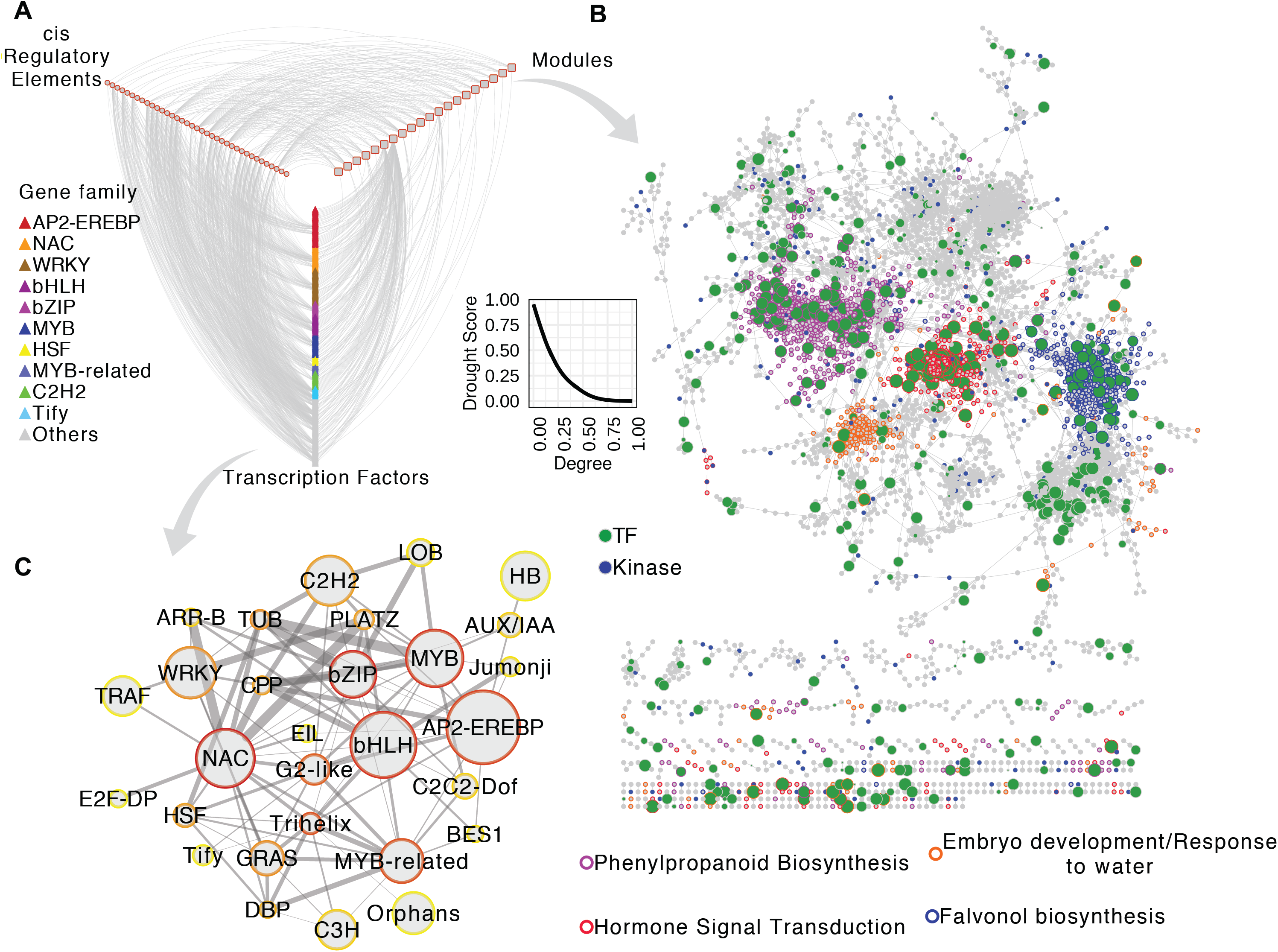
Functional characterization of predicted drought tolerance transcription factors. A) A subset of modules with highest feature importance scores from the drought classifier were connected to cis-regulatory elements (CREs; predicted by *de novo* analysis) found enriched within them, as well as to their predicted regulators (TFs). The regulators were in turn connected to the CREs based on enrichment analysis (FDR corrected hypergeometric test *p* value < 0.01). This interconnected network with three node types (modules, CREs, TFs) was visualized in Cytoscape. Modules are indicated in rounded rectangles, CREs in ellipses and TFs in triangles colored according to the family membership. Inset shows relationships between drought score and network degrees of TFs. B) The drought modules consist of a total of ∼6000 genes. The network shows top 5% edges induced between them. Every grey circle is a functional gene, green circle is a TF and blue circle is a kinase. Size of green circles is proportional to the drought score. Border of nodes that co-occur in the same module are given the same color, and the module function is indicated in the key below. C) TFs were connected to each other based on mutual information between their network profiles to create a global TF-TF network. In the global TF-TF network, the sum of cross-family edge-scores were summarized as *Z* scores. Pairs of TF families with high *Z* scores were visualized as a graph. Each ellipse represents a TF family, with node size proportional to the total number of members within the family and border color set along a yellow to red gradient indicating to the total number of connections with TFs in other families. Colors closer to dark red indicate larger number of connections and colors closer to yellow indicate fewer connections. Edge thickness is proportional to the *Z* score of connection between the two families linked.

It is important to note that predicted DS did not simply reflect on ‘hubness’ of TFs in the GRN (**Fig. 6A inset**). Instead, the predicted DT TFs appear to be involved in the regulation of a small number of key drought modules. These drought modules comprise a total of 6968 genes which form core communities enriched in several stress response pathways and biological processes **(Fig. 6B)**. Interestingly, ‘hormonal signal transduction’ and related pathways such as ‘phenylpropanoid biosynthesis’ and ‘jasmonic acid biosynthesis’ were found most strongly enriched in this network. Because most TFs with large DS in our predictions arose in land plants (linked to vascular development), this functional enrichment pattern is in strong agreement with a recent study that showed evolution of abscisic acid and salicylic acid pathways, along with jasmonate signaling pathways, in land plants (Wang et al., 2015). Interestingly, the most prominent *de novo* predicted AS-CREs in this network are also related to the abscisic acid response complex ABRE3HVA22 (Shen et al., 1996) and the vascular-specific motif ACIIPVPAL2 (Hatton et al., 1995), along with the light responsive GT-1 motif (Lam and Chua, 1990), the anerobic-responsive motif GCBP2ZMGAPC4 (Geffers et al., 2000) and the dehydration responsive DREB1A motif (Maruyama et al., 2004) (**Supplemental Data S4)**. Other relevant GO biological process terms such as ‘response to water’, ‘response to abscisic acid stimulus’, ‘cellulose biosynthesis’, ‘flavonol biosynthesis’ and ‘trehalose biosynthesis’ were also correctly recovered in this drought network.

As mentioned previously, TFs at the top of our rankings are enriched in known stress related gene families. We next investigated the extent to which TFs liaise with other TFs in different families by estimating mutual information between their network connectivity profiles **(Fig. S4;** see *Supplemental methods*). We observed that the members of AP2-EREBP, bHLH, NAC, MYB and bZIP families have the largest number of cross-family interactions **(Fig 6C)**. Surprisingly, the seemingly under-studied CPP (cysteine-rich polycomb-like protein) family showed strong connections to these hub families, suggesting their important role in drought response. A previous study reported on the classification of CPP genes from multiple plant species into two distinct groups, based on their protein domain features (Lu et al., 2013). The authors suggested that TFs in these two groups could likely be independently involved in distinct cellular functions. We confirm this hypothesis, and suggest that group 1 members of the CPP family are possibly involved in stress response pathways; 4 of the 5 members of group 1 were predicted positive by our drought classifier, while all members of group 2 were predicted negative. The one mis-classified TF (LOC_Os04g09560) from group 1 could likely be due to a different domain architecture compared to rest of the members of the same group (Lu et al., 2013).

Overall, network analysis showed that TFs predicted to be involved in DT mechanisms are more likely to bind to CREs commonly implicated under abiotic stress, functionally cooperate with other TFs from same and other families, and function in regulation of network communities involved in hormonal signaling.

### DroughtApp allows functional characterization of rice genes

We developed a user-friendly webserver called DroughtApp with the intention to provide an easy interactive access to the consensus GRN and drought predictions we described here. The DroughtApp is built using R/Shiny framework and allows users to browse the network neighborhood of genes of interest. We chose the rice transcription factor *bHLH148* (LOC_Os03g53020) TF to demonstrate how the DroughtApp could be integrated in systems biology projects to generate new testable hypothesis. It shows that *bHLH148* was strongly predicted for its association with DT, and its predicted target genes in the consensus GRN are other TFs from the WRKY and AP2-EREBP families **(Fig. S5)**. To experimentally validate these predictions, we first verified the association of *bHLH148* with drought stress at different stages using a homozygous loss-of-function knockout mutant line designated as ‘bhlh148’ **(S6 A-C)**. We tested the drought stress response of *bhlh148* plants under controlled drought stress. Under well-watered condition, there were no significant phenotypic difference between the mutant and WT plants. But under controlled drought stress treatment at 40% field capacity (FC), the mutant plants showed higher sensitivity with leaves rolled and collapsed compared to the WT plants **(Fig. 7A)**. Under drought, the *bhlh148* mutant plants showed significant reduction in net photosynthetic rate, instantaneous water use efficiency (WUEi), efficiency of Photosystem II measured in light adapted leaves (Fv′/Fm′), relative water content (RWC) and the above ground biomass compared to WT **(Fig 7B-F)**. Further, yield parameters for drought stress response quantified by number of panicles **(Fig. 8A)**, number of spikelets **(Fig. 8B)**, percent spikelet sterility **(Fig. 8C)** and grain yield per panicle **(Fig. 8D)** testify that bHLH148 is involved in grain yield under drought stress (see *Additional notes*).

**Figure 7:**
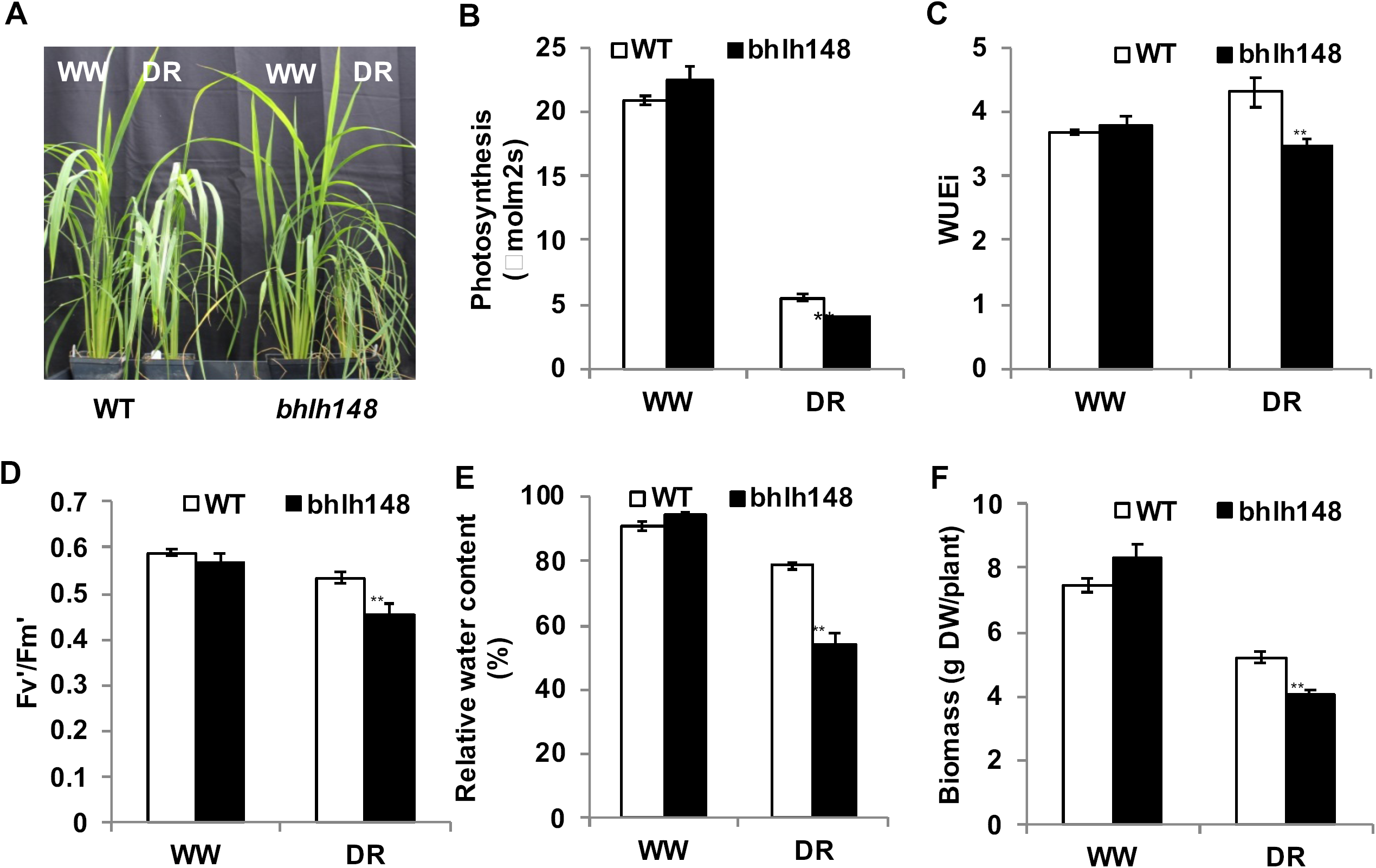
Drought induced expression of *bHLH148*. A) Increased sensitivity of *bhlh148* mutant plants under controlled drought stress conditions. Forty-five-day old plants were maintained at 100% (well-watered – WW) and 40% (drought – DR) FC (field capacity) for 10 days by a gravimetric approach and performance was measured at the end of stress period. B-F) Phenotype of the WT and *bhlh148* mutant plants under drought stress. B), Assimilation rate C), instantaneous water use efficiency (WUEi) D), efficiency of Photosystem II in light adapted leaves E), and relative water content (RWC) F) and above ground biomass (dry weight). Gas exchange measurements were taken using portable photosynthesis system LI-6400XT at CO_2_ concentration of 370 μmol/mol and light intensity of 1000 μmol/m^2^/s. The data are the means ± s.e. (n=10) and significance using *t-*test (\*\**P* ≤ 0.01). K-N)

**Figure 8:**
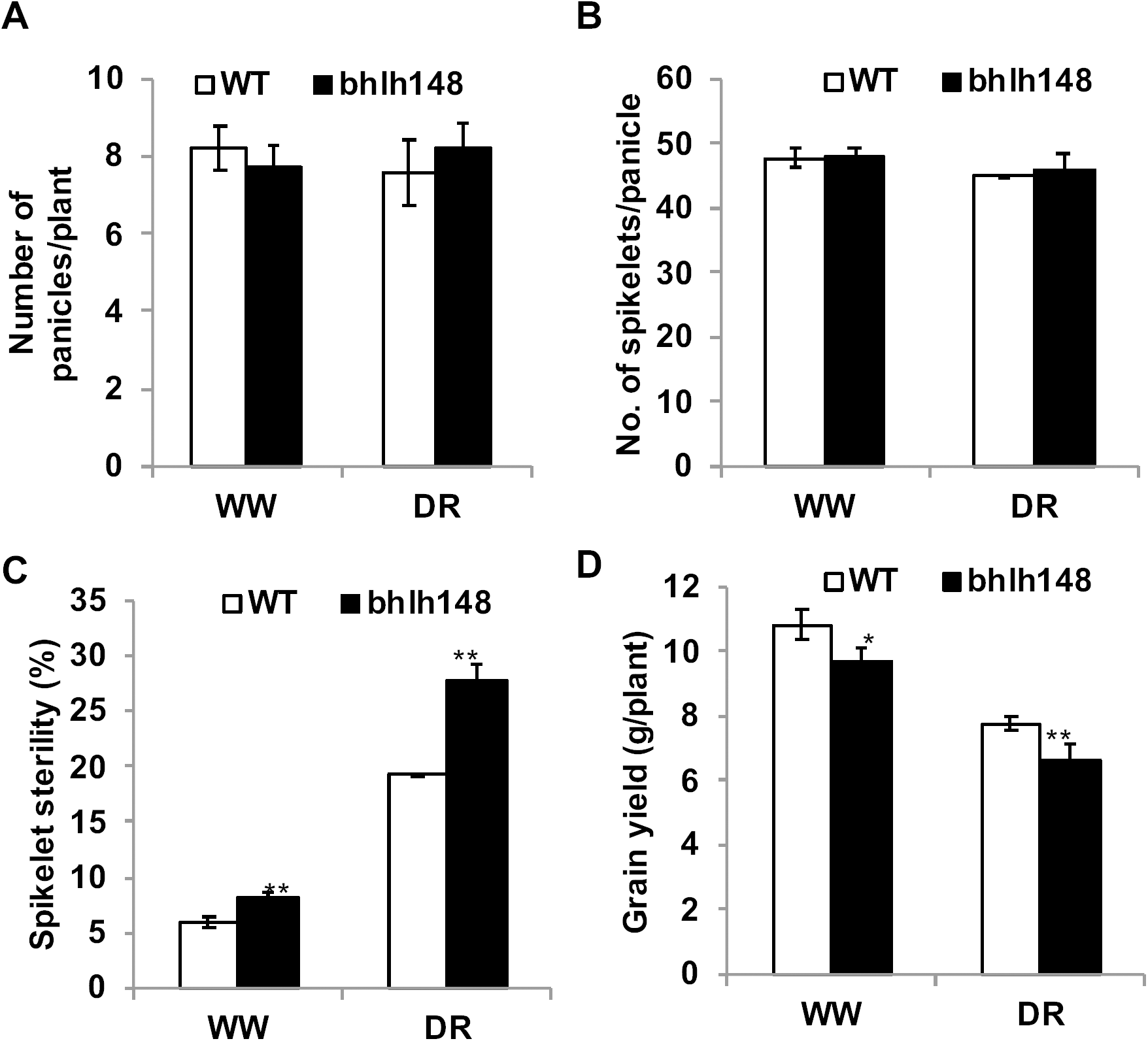
Reduced grain yield of *bhlh148* plants under well-watered as well as drought stress conditions. Drought stress was applied by withholding irrigation at R3 stage for 4-8 days until the leaves roll and wilt followed by re-watering and maintaining under well-watered condition until physiological maturity. Yield components were measured under well-watered and drought stress conditions at physiologically maturity. Number of panicles, B) number of spikelets, C) percent spikelet sterility and, D) grain yield. The data are means ± s.e. (n=6) and significance using *t-*test (\**P* ≤ 0.05 and \*\**P* ≤ 0.01).

To verify whether *bHLH148* targets the WRKY and AP2/EREBP family of TFs as predicted by the DroughtApp, we performed gene expression profiling of bhlh148 and WT plants under well-watered and controlled drought stress conditions using RNA-sequencing (see *Supplemental Methods*). Leaf tissue from plants maintained at 100% and 40% field capacity for 10 days, were used as well-watered and controlled drought stress samples, respectively. Analyses of differential expression was performed to identify genes that 1) responded to the knockout, 2) responded to drought in WT plants, and 3) respond specifically to the interaction of mutant with drought (subtracting the baseline effect of drought from mutant) (**Supplemental Data S7)**. Subsequently, functional enrichment tests using MapMan terms were performed using fold change values as a parameter to evaluate significantly up- and down-regulated pathways (Kim and Volsky, 2005). These analyses showed that transcripts annotated to ‘regulation of AP2/EREBP element binding protein family’ and ‘regulation of WRKY domain TF family’ were strongly downregulated, specifically in the drought treated mutant plants **(Fig. 9A; Supplemental Table 2)**. We found that 67% (55/81) of TFs predicted as targets of *bHLH148* were significantly differentially expressed in the WT plants exposed to drought (*q* < 0.01), confirming their predicted high DS (**Fig. S7**).

**Figure 9.**
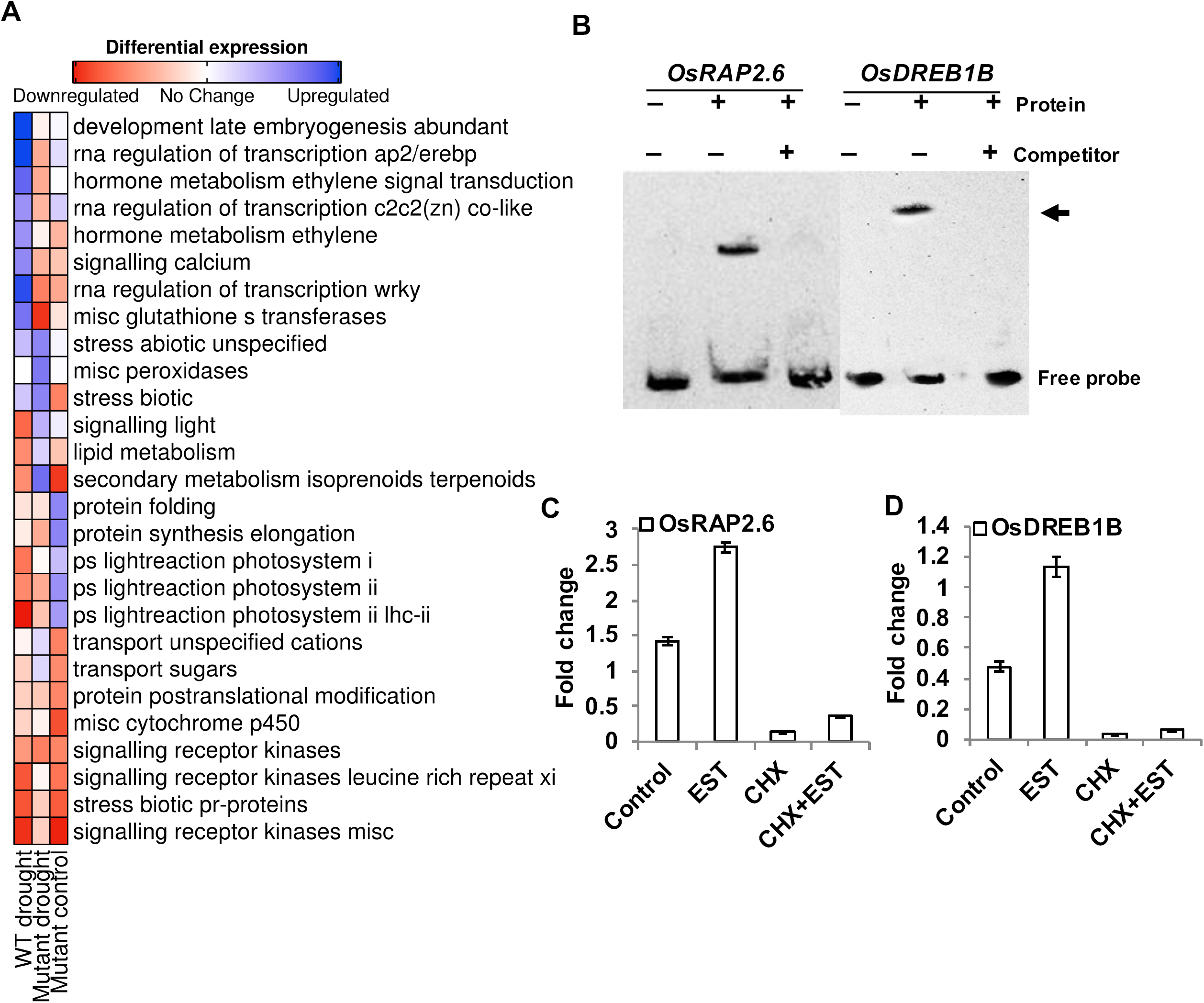
Experimental validation of DroughtApp predictions. A) Heatmap summarizing results from differential gene expression analysis of the rice bhlh148 mutant exposed to drought. The heatmap shows the average differential expression of gene transcripts annotated to various pathways listed in the rice MapMan database. The color gradient indicates mean fold change (summarized as *Z* scores) of the pathway listed in the row and sample in the column. The color gradient represents up and downregulation, as indicated in the color key above. B) Electrophoretic mobility shift assay (EMSA) was performed with bHLH148 protein and biotin labeled promoter elements of potential bHLH148 regulated genes. bHLH148-6xHis recombinant protein was incubated with promoter elements at room temperature for 20 min. For competition analysis, the binding reaction was incubated for 10 min on ice before adding 100-fold excess of unlabeled promoter elements followed by incubation at room temperature for 20 min. The samples were subjected to EMSA by PAGE and subsequent chemiluminescence detection. + and − indicate the presence and absence of the respective component in the binding reaction. The labeled “free probe” and DNA-protein complex “bound probe” positions are indicated by arrows. C-D) Direct activation of *OsRAP2*.*6* and *OsDREB1B* by bHLH148. Rice protoplasts were transfected with a bHLH148-HER fusion construct driven by the CaMV35S promoter. Transfected protoplasts were treated with estradiol (EST), cycloheximide (CHX), or EST and CHX together. The expression levels of *OsRAP2*.*6* and *OsDREB1B* in control and treated protoplast was analyzed by qPCR and shown for (C) *RAP2*.*6* and (D) *OsDREB1B*. Each data point are mean values ± s.e. of three biological replicates.

We next tested whether bHLH148 can directly bind to the E-box elements on the promoters of a few differentially expressed AP2/ERF genes that were also predicted as targets of *bHLH148* by the DroughtApp (**Supplemental Data S8)**. To do this, we performed an electrophoretic mobility shift assay (EMSA) and confirmed that bHLH148 binds to the promoters of OsRAP2.6 (LOC_Os08g36920) and OsDREB1B (LOC_Os09g35010) genes **(Fig. 9B;** see *Additional notes*). To further verify whether bHLH148 can directly activate the expression of AP2/ERF genes that were identified by EMSA, we used the steroid receptor-based inducible system, and confirmed that bHLH148 directly activates expression of OsRAP2.6 **(Fig. 9C)**, while activation of OsDREB1B by bHLH148 requires additional factors **(Fig. 9D)**. Among these two AP2/ERF TFs, the role of OsDREB1B in imparting drought stress tolerance to rice plants has been shown through activation of several stress responsive genes (Ito et al., 2006). The role of OsRAP2.6 (ERF101) in regulation of drought in reproductive tissues has been recently revealed (Jin et al., 2018), which also supports the observed grain-yield phenotype of bhlh148.

## Conclusion

Our survey of the literature and mining of phenotype databases show that currently only ∼2% (1098 at the time of this study) of all known rice genes have been linked to various abiotic stresses experimentally, but more than 15% of these 1098 stress genes are TFs linked with drought or water deficit related responses. This suggests that genetic selection of favorable alleles of the stress inducible TFs has been widely and inadvertently used as a tool to improve/select for drought tolerance. We leveraged on regulatory network patterns of these experimentally validated examples of drought regulators to train machine learning models for genome-wide prediction of TFs and associated physiological pathways involved in various drought tolerance (DT) mechanisms of rice. Unlike traditional coexpression analysis, our supervised approach allowed us to rank each TF in the rice genome according to its predicted association to DT, and these rankings could be objectively tested. We anticipate that our predictions will be a valuable resource for exploring the transcriptional regulatory code of plant responses to drought stress.

The strategy described ultimately led to the characterization of TFs most likely to be involved in DT mechanisms. A strong enrichment of intron-poor TFs among the top of the genome-wide ranking suggests that drought regulators are more likely to be rapidly regulated in response to drought stress (Jeffares et al., 2008). Widespread upstream regulation of these TFs was also suggested by the large presence of *de novo* predicted stress-relevant *cis* regulatory elements within their promoters relative to other TFs. The strongest enrichment of their predicted target genes was found with modules involved in hormone-mediated signaling, along with the phenylpropanoid pathway and other smaller pathways that depend on it **(Fig. 6B)**. It is important to note that most of the top ranked TFs in our analysis emerged in land plants **(Fig. 4F)**. Thus, the functional enrichment patterns indicate that the phenylpropanoid pathway, which is also implicated in lignin biosynthesis (Fraser Cm Fau - Chapple and Chapple), played an important role in adaption of plants to water limiting environments, as also suggested in recent reports (Wang et al., 2015; Ahammed et al., 2016; Verma et al., 2016). Even the most strongly enriched CREs involved in regulation of in the drought modules agree with these functional roles of predicted drought response regulators. The rankings estimated here provide a primer to experimentally explore functional features of drought TFs by recording their phenotypes conditioned on drought stress. The network-based machine learning approach presented here, in conjunction with resources like the KitaakeX Mutant Database (Li et al., 2017), can support targeted screens to narrow down the search for TFs involved in specific physiological, morphological and biochemical phenotypes of drought response. This will in turn enable classification for a specific phenotype in future studies.

Nested cross-validation tests suggested that models trained using network connectivity patterns as features are generally more accurate and robust to variation in training labels **(Fig. 5G)**. The approach we present here can potentially be applied across transcriptomes within many biological contexts for which enough training labels are also available. However, the generalizability of trained models will depend upon the quality of training examples, standard of validation data and feature engineering. The observed drop in accuracy of the model trained with integrated genomic and network features was expected due to the increase in model complexity. Nevertheless, it also suggests that this technique of integrating different data-types is feasible, and opens new avenues for development of more mechanistically informed models. Integration of the transcriptional-level regulatory code of drought response we present here with other diverse sources of information – representing different layers of TF mediated gene regulation – into a single model predictive of drought response genetics will allow candidate gene selection in a truly holistic manner. These new datasets should be inclusive of tissue-specific network models, epigenetic profiles, frequency of alternative splicing, post-transcriptional regulation by microRNAs and post-translational modifications (PTM) such as phosphorylation. Some excellent resources, such as the Plant PTM Viewer (Willems et al., 2019) and the database of phospho-sites in plants (Cheng et al., 2014) currently allow such data mining for a few plant TFs. Perhaps, such an integration could also help achieve a better classification of functional alleles in *indica* and *japonica* sub-types of rice, which remains a limitation of our study.

## Methods

### Creating the consensus gene regulatory network

A set of 35 Affymetrix microarray datasets comprising of 266 individual samples pertaining to gene expression profiling of rice plants under the context of abiotic stress were identified in GEO (**Supplemental Data S9**). The raw data was downloaded, individually normalized and processed into an integrated expression matrix as previously described (Krishnan et al., 2017). A comprehensive list of 2304 known rice genes annotated as TFs in several public databases was curated over years (Yilmaz et al., 2009; Jung et al., 2010; Priya and Jain, 2013; Jin et al., 2014). This list of TFs, along with the normalized gene expression matrix was supplied to five reverse-engineering algorithms. ARACNE was downloaded from the web link in the original publication. GENIE3 (Huynh-Thu et al., 2010) and CLR (Faith et al., 2007) runs were performed using the R package minet (Meyer et al., 2008). Each of these algorithms required calculation of mutual information (MI) between every possible TF-gene pair. Bootstrapping was avoided because genome-wide calculations of MI in rice is computationally intensive. Top 500,000 edges were selected from the output of each of these three algorithms and from the two correlation-based methods. The union of all edges from all methods was used to create an edge matrix *E*, with edges *i* in rows of *E* and algorithms *j* in columns of *E*. Each cell in the *E*_*ij*_ was populated by the rank given to *i* by *j*. Missing edges were substituted with the lowest rank of that column plus one (Marbach et al., 2012). The average rank for each row was then computed and ranked. Hence, edges with small values indicated greater confidence by all five methods. Top 500,000 edges from this aggregate were selected as the consensus gene regulatory network (GRN) of rice. Estimation of coregulation amongst gene-pairs and network modules were identified using the technique described previously (Vermeirssen et al., 2014) (see *supplemental methods*).

### Creation of validation networks

The Position Weight Matrices (PWM) of ∼588 rice TFs listed in the CIS-BP database (Weirauch et al., 2014) were obtained in April 2019. PWMs indicate DNA sequence preferences of TFs and can be used to infer DNA motifs in the promoter regions of functional genes. The 1000bp upstream promoters were scanned for at least one or more occurrence of the PWM motifs using the FIMO tool in the MEME suite (Bailey et al., 2015). Motifs that were found in more than 50% of all the genes were treated as ‘constitutive elements’ and removed. Genes harboring all the remaining motifs with a *p*-value < 1E-10 were linked to the corresponding TFs and used for evaluations. The functional evaluation network was created by using evidence of functional relationships between TFs and putative target genes co-annotated in the rice biological process (BP) ontologies and MapMan pathways. Only those annotation labels consisting of less than 200 genes were chosen for this. We assumed that TF within each of these specific BP terms and pathways are more likely to be direct regulators of all other genes within the same term or pathway, and at least these links should be predicted with greater confidence even if they are indirect. Excluding large BPs and pathways, we ensured that minimally related genes (in processes such as ‘translation’, ‘DNA repair’, ‘signal transduction’ etc.) did not become part of the validation network. A total of 242 TFs were found co-annotated with 4670 functional genes in GO BP, and 1520 TFs were found co-annotated with 4021 functional genes in the MapMan database. Both these validation networks were used to calculate the precision and recall statistics and the *F-*score (see *Additional notes*).

### Generating training labels for machine learning

To identify drought positive labels, the gene keyword file from the funcricegenes server was obtained https://funricegenes.github.io/ in May 2019. Gene lists available in the Oryzabase database was obtained from https://shigen.nig.ac.jp/rice/oryzabase/download/gene on the same day. The rice mutant database were obtained from the published article (Zhang et al., 2006). Using a word cloud analysis (not shown), most prominent keywords in these databases were visualized. Genes linked with keywords related to abiotic stress such as “drought”, “water-deficit”, “salt”, “cold”, “heat”, “temperature” and “disease” were then extracted. The retrieved locus IDs and publication records of genes were manually scanned for consistency by expert stress biologists, and TFs linked with drought (and related keywords) were labeled as positives. Note that OsbHLH148 was originally present in our dataset as a drought positive TF (Seo et al., 2011), but it was removed from the positive list prior to training the models as a hidden example on which wet-lab experiments were performed later. From the remaining TFs, we listed negatives examples as those that were not positive for any abiotic stress in database mining, since many genes are multi-stress responsive. Also, those TFs that did not differentially expressed in reanalysis of seven published gene expression datasets covering drought stress responses in various organs and tissues of rice plants across multiple genotypes were also counted as drought negatives. In addition to this, the rice stress TF database was downloaded (Priya and Jain, 2013) from http://www.nipgr.ac.in/RiceSRTFDB.html and TFs not listed as responsive to drought and salt in this database was also included as negative TFs. Altogether, we created a pool of 752 TFs that are most likely not regulators of drought stress responses. To build an unbiased model, we randomly selected ‘hold-out set’ of 422 TFs (∼ 50% of the combined list of all positive and negative TFs). This hold-out set was later used to evaluate the performance of the final model. The remaining 50% of labeled TFs were used in the training dataset for the network-based classifier.

### Network-based classifier

The modular core of the consensus GRN we inferred was structured as a matrix *G*, with each entry in *G*_*ij*_ corresponding to the Jaccard coefficient (JC) of TF *i* in row with module *j* in the column. 590 modules that were found to be functionally enriched, coregulated by the same sets of abiotic stress CREs or preserved in an independent coexpression network were considered biologically relevant and used in *G*. The subset of *G* with JC values of training labels was supplied to a linear kernel support vector machine (SVM) classifier. The vector of JC values of each labeled TF across all modules in *G* represented its feature vector. The objective of an SVM function is to identify the best hyperplane that separates two classes of training data (drought positive and negative TFs) using their feature vectors. The width of the margin that separates the two classes was controlled by optimizing the classification trade-off parameter (*C*; a penalty for a miss-classified example). An optimal *C* was chosen by testing a range of values from 0.001 to 10 in increments of 0.1 and five-fold cross validation tests. Classifier training runs were performed using the libSVM package (Chang and Lin, 2011). Cross validation splits and performance evaluation was performed using the ROCR package in R (Sing et al., 2005). The distance from the hyperplane for each TF returned from the final SVM run was averaged over four values from five-fold cross validation runs. The entire range of these average distances were scaled to the range 0 to 1. The resulting value of each TF was treated as its drought score (DS).

### Feature engineering for the integrated model

#### TF-DNA binding sites

The binding motifs of rice TFs was obtained from the CIS-BP database (Weirauch et al., 2014). These motifs were first matched with *de novo* predicted motifs (from FIRE) using the TomTom tool in MEME (Bailey et al., 2015). Matching motifs with a *q* value < 0.1 were then removed from the CIS-BP group of motifs, as FIRE predicted motifs were considered stress-specific. TFs were linked to FIRE motifs by overlap analysis with predicted targets of TFs in the GRN (hypergeometric tests *q* value < 0.01). All TF-motif links from CIS-BP and FIRE analyses were then combined to create a non-redundant set of putative CREs, represented as a matrix *C* with TFs *i* in rows and motifs *j* in columns. Each cell *C*_*ij*_ was populated with 1 if a link between row TF and column motif was observed, 0 otherwise.

#### TF families

TF family annotations were downloaded from the Plant TF database (http://plntfdb.bio.uni-potsdam.de/v3.0/downloads.php?sp_id=OSAJ). Gene-family relationships were represented as a matrix *F* with TF *i* in the row and family name *j* in the column. Each cell in *F*_*ij*_ was populated with 1 if the *i* is a member of *j*, 0 otherwise.

#### Response to hormones

The dataset GSE37557 was downloaded from GEO and differential expression quantified using method previously described (Krishnan et al., 2017). The cells of a matrix *H* with TFs *i* in rows and six hormone treatments *j* in columns was filled with 1 if *i* had a positive fold change in the treatment represented by *j*, 0 otherwise.

#### Network degrees

Outdegrees of all TFs from the TF-TF mutual information network was divided into quantiles, and a TF was assigned to one of the four quantiles. The matrix *D*_*ij*_ with TFs *i* in rows and each of the quantile *j* in columns was accordingly populated with either 1 or 0.

*Gene age* was obtained from the gene feature file obtained from rice pan genome server (http://cgm.sjtu.edu.cn/3kricedb/data/GeneFeature.txt). In this feature file, the age column had 13 NCBI taxonomic classes labeled as PS1 to PS13 (Phylostratum 1-13). The matrix *A* with TFs *i* in rows and each of the 13 phylostrata in the column *j* was filled with 1 if *i* was found assigned to the age group represented in *j*, 0 otherwise.

#### Structural features

Number of protein domains per TF was obtained from the ‘all.interpro’ file available in the download section of the rice genome annotation project website (http://rice.plantbiology.msu.edu/). TFs were grouped according to the number of interpro domain annotations. The matrix *P* with TFs *i* in rows and five groups in columns *j* was filled with 1 if *i* was found to have that many numbers of protein domain(s) represented in *j* (e.g. TFs in group 1 have 1 domain, group 2 have 2 domains, and so forth). Number of introns per TF was calculated from the GFF file of rice reference genome. The matrix *I* with TFs *i* in rows and number of introns *j* was populated with 1 if *i* had that many introns indicated in *j*, 0 otherwise.

Finally, the matrices *C, H, F, A, D, P* and *I* were integrated with the GRN matrix *G* to create the integrated feature matrix for 2160 TFs and 4597 features. All missing values in the integrated matrix were substituted with 0.

### Analysis of RNA-seq data and estimation of differential expression

Raw fastq files of individual samples from all external datasets were downloaded from the SRA. The Nipponbare RefSeq (MSU version 7) was obtained from the rice genome annotation project website (Kawahara et al., 2013). The barley and sorghum genomes and annotations were downloaded from the Phytozome web portal (Goodstein et al., 2012). The following procedure was applied across all RNA-seq samples, including samples from mutant experiments generated in the study described here. Reads were mapped to the respective reference genomes using STAR version 2.7 (Dobin et al., 2013). The bam files obtained from STAR runs we sorted using samtools and used as input to the HTseq software version 0.11.2 (Anders et al., 2015) with its default parameters for counting reads per gene per sample. Count of reads obtained from HTseq runs were then integrated as a count matrix (one for each experiment) with columns representing individual samples and rows representing genes, and each cell of the matrix presenting raw counts of the gene in the corresponding sample. The count of each gene in the count matrix was first scaled by its length to give reads per kilobase (RPK). The sum of all RPK values per sample divided by 1 million gave us a scaling factor, and dividing each RPK value by the scaling factor computed gene expression as transcripts per million (TPM) units. The effective gene length to be used in calculations of RPK values was computed as the sum of non-overlapping exon lengths using the genomic features package in R (Lawrence et al., 2013). The GFF3 files of all genomes were converted to GTF format using GFF utilities (gffread) of the cufflinks software (Trapnell et al., 2010). The resulting GTF file was used as input to genomic features for effective gene length calculation. Note that the rice GFF3 file on rice MSU reference has mis-annotations of ∼1000 gene isoforms, which hampered gene length calculations. Conversion of GFF3 to GTF ensured proper grouping of individual transcripts to parent gene ID. For test of differential expression, the raw count data was normalized using voom (Law et al., 2014) and differential expression of genes between control and treatment samples was estimated from linear models using the limma package in R (Ritchie et al., 2015). Differential expression from microarray datasets **(Fig. 4A-E)** was estimated using the procedures described previously (Ambavaram et al., 2014).

### Controlled drought stress at vegetative stage and physiological measurements in rice

To test the drought stress response of mutant plants at the vegetative stage, we applied controlled drought stress on 45-d-old plants using a gravimetric approach. One-week old equal sized individual seedlings were transplanted into 4 square inch plastic pots filled with Redi-earth potting mix of known weight and water holding capacity. Thirty-five days after transplanting, controlled drought stress (DR) was initiated on 10 pots and monitored gravimetrically. The soil water content was brought down to 40% FC over a period of 3 to 4 d and plants were maintained at that level for 10 d by weighing the pots daily at a fixed time of the day and replenishing the water lost through evapotranspiration. Another 10 pots were maintained at 100% FC and treated as well-watered (WW) condition (Ramegowda et al., 2014). At the end of the stress period, gas exchange and light adapted fluorescence measurements (Fv′/Fm′) were taken on the 2^nd^ fully expanded leaves from the top, using a portable photosynthesis meter, LI-6400XT (LI-COR Inc., NE, USA) at CO_2_ concentration of 370 μmolmol^-1^, light intensity of 1000 μmolm^-^2s^-1^ and RH of 55-60%. Instantaneous water use efficiency (WUEi) was calculated using net photosynthetic rate (A) and transpiration rate (T) as WUEi = (A/T). Leaf RWC was measured as described (Barr and Weatherley, 1962) in the leaves used for gas exchange measurements. The leaf fragments of same length were excised and fresh weight (FW) measured immediately. Leaf fragments were hydrated to full turgidity by floating them on deionized water for 6 h, then blotted on paper towel and the fully turgid weight (TW) taken. The leaf samples were then oven dried at 80°C for 72 h and weighed to determine dry weight (DW). The percent RWC was calculated as RWC (%) = (FW - DW)/(TW - DW) x 100. To determine biomass, shoots were harvested, oven dried at 80°C for 72 h and weighed.

### Grain yield analysis under reproductive drought in rice

The effect of drought stress on grain yield of the rice genotypes was tested by applying drought stress to plants at R3 stage (Counce et al., 2000). Individual plants in 4 square inch plastic pots were grown at well-watered conditions until R3 stage. Drought stress was applied by withholding water at R3 stage for 4 to 8 d until all of the leaves wilted followed by re-watering. Panicles exposed to drought stress during the 4 to 8 d window were marked and used for yield component analysis. A set of well-watered plants were also maintained as controls. Plants were further grown in well-watered condition until physiological maturity. Drought exposed panicles were harvested and number of filled and unfilled spikelets counted to determine spikelet sterility (%). The filled spikelets were dried at 37°C for 5 d and weighed to determine grain yield/plant.

### Electrophoretic mobility shift assay (EMSA)

The total RNA isolated from drought stressed rice plants was used to amplify full-length cDNA encoding bHLH148 and cloned into pET28(a) vector at *Bam*HI and *Eco*RI sites. The bHLH148-6xHis recombinant fusion protein expression was induced with 1 mM IPTG for 4 h and purified using Ni-NTA resin, and the identity of the purified protein was confirmed by western blotting (data not shown) using the His-tag antibody. The binding reaction and EMSA were carried out using a standard protocol according to the manufacturer’s instructions (LightShift Chemiluminescent EMSA Kit). Promoter sequences (2 kb upstream of transcription start site) of AP2/ERF TFs were identified using PlantPAN database (http://plantpan.mbc.nctu.edu.tw/) (Chang et al., 2008) and searched for the presence of E-box elements in the PLACE database (http://www.dna.affrc.go.jp/PLACE/) (Higo et al., 1999). Specific sets of primers were used to amplify 200 bp E-box flanking regions of each of the putative bHLH148-regulated gene promoters using rice genomic DNA as a template. The amplified promoter fragments were biotin labelled at the 3′ end using the Biotin 3′ End DNA Labelling Kit (Pierce). The binding reactions were carried out in a buffer containing 10 mM Tris (pH 7.5), 50 mM KCl, 1 mM dithiothreitol, 2.5% glycerol, 5 mM MgCl, 0.05% Nonidet P-40, and 50 ng/µl of poly(dI-dC). For competition analysis, the binding reactions were incubated for 10 min on ice before adding 100-fold excess of unlabelled competitor DNA, and the reaction mixture was further incubated for 20 min at room temperature before loading onto a 5% native polyacrylamide gel. The resolved DNA-protein complexes were electro-blotted onto nylon membranes and subsequently detected using the chemiluminescence detection kit.

### Steroid-inducible system for testing direct activation of genes by bHLH148

The bHLH148-HER expression construct was generated by ligating the PCR-amplified full-length cDNA of bHLH148 at the *Kpn*I site fused with the regulatory region of HER at the C terminus between the CaMV 35S promoter and the NOS terminator in pUC19 vector. The construct was transfected into rice protoplasts by electroporation and incubated with 2 μM estradiol for 6 h to release cytoplasmic bound bHLH148. For the control reactions, the same concentration of ethanol used to dissolve estradiol was used. To inhibit new protein synthesis, protoplasts were treated with cycloheximide (2 μM) for 30 min before addition of estradiol. Total RNA was isolated from the treated protoplasts and used for qPCR analysis. The data presented are the averages of three biological replicates.

## Supplemental Datasets

**Supplemental Data S1:** Top 500,000 edges inferred by the ensemble and their aggregate.

**Supplemental Data S2:** Gene-module memberships

**Supplemental Data S3:** Module function annotations

**Supplemental Data S4:** Module CREs annotations

**Supplemental Data S5:** Drought Scores

**Supplemental Data S6:** Feature importance scores

**Supplemental Data S7:** Differential expression test results from all three analyses

**Supplemental Data S8:** Predicted targets of bHLH148 from DroughtApp

**Supplemental Data S9:** GEO datasets

## Data availability

All RNA-seq datasets published with this study are deposited to the NCBI repositories and can be accessed through GEO accession GSE65024.

## Acknowledgements

This study was supported from the NSF-MCB award 1716844: Systems genetics analysis of photosynthetic carbon metabolism in rice: and the NSF-EPSCoR award 1826836: RII Track-2 FEC: Systems genetics studies on rice genomes for analysis of grain yield and quality under heat stress. This research is also supported by the Arkansas High Performance Computing Center which is funded through multiple National Science Foundation grants and the Arkansas Economic Development Commission. The authors would like to thank Arjun Krishnan of the University of Michigan, and members of the Pereira lab for relevant discussions on the topic. The authors would also like to thank the original contributors of all publicly available datasets used in this study.

